# The C-terminal acid phosphatase module of the RNase HI enzyme RnhC controls rifampicin sensitivity and light-dependent colony pigmentation of *Mycobacterium smegmatis*

**DOI:** 10.1101/2022.11.09.515703

**Authors:** Pierre Dupuy, Michael S Glickman

## Abstract

RNase H enzymes participate in various processes that require processing of RNA:DNA hybrids, including DNA replication, transcription, and ribonucleotide excision from DNA. Mycobacteria encode multiple RNase H enzymes and prior data indicates that RNase HI activity is essential for mycobacterial viability. However, the additional roles of mycobacterial RNase Hs are unknown, including whether RNase HII (RnhB and RnhD) excises chromosomal ribonucleotides misincorporated during DNA replication and whether individual RNase HI enzymes (RnhA and RnhC) mediate additional phenotypes. We find that loss of RNase HII activity in *M. smegmatis* (through combined deletion of *rnhB/rnhD*) or individual RNase HI enzymes, does not affect growth, hydroxyurea sensitivity, or mutagenesis, whereas overexpression of either RNase HII severely compromises bacterial viability. We also show that deletion of *rnhC*, which encodes a protein with an N terminal RNase HI domain and a C terminal acid phosphatase domain, confers sensitivity to rifampicin and oxidative stress as well as loss of light induced carotenoid pigmentation. These phenotypes are due to loss of the activity of the C terminal acid phosphatase domain rather than the RNase HI activity, suggesting that the acid phosphatase activity may confer rifampicin resistance through the antioxidant properties of carotenoid pigment production.

## INTRODUCTION

The basic unit of RNA, ribonucleotide triphosphates (rNTPs), are more abundant than the basic unit of DNA, deoxyribonucleoside triphosphates (dNTPs), in the nucleotide pool of eukaryotic and prokaryotic cells (1, 2). rNTPs differ from dNTPs by the presence of a hydroxyl group on the 2’ carbon of the ribose, which physically clashes with a bulky residue of replicative DNA polymerases, termed the steric gate, thereby limiting ribonucleotide incorporation into chromosomal DNA during replication (3–5). Despite the steric gate, replicative polymerases still incorporate ribonucleotides at an estimated rate of 1/1000 bases replicated (2). Genome replication also incorporates ribonucleotides into DNA during lagging strand replication in the form of RNA primers used for Okazaki fragments (6–8). Finally, R-loops, formed when the nascent RNA anneals with the template DNA strand during transcription (9–11) is yet another form of the intermingling of RNA with DNA. Cellular processing of these RNA:DNA hybrids is essential for core cellular functions, including DNA replication, transcription, and repair. Failure of properly process RNA:DNA hybrids induces genome instability by increasing mutations and recombination (12–15).

To prevent deleterious effects of persistent genomic ribonucleotides, organisms encode RNase H enzymes which incise the RNA strand of RNA:DNA hybrid duplexes (16–18). RNase H enzymes are widely distributed in eukaryotes, prokaryotes, and retroviruses (19). They are classified into two main classes: RNase HI and RNase HII (16–18). RNase HI acts on a tract of at least four ribonucleotides but cannot incise a single ribonucleotide embedded in duplex DNA, whereas RNase HII can incise a single embedded ribonucleotide. RNase HI enzymes are involved in the degradation of R-loops and unprocessed RNA primers from Okazaki fragments, whereas RNase HII initiates the ribonucleotide excision repair (RER) pathway, removing single ribonucleotides from genomic DNA (16–18).

*Mycobacterium tuberculosis* (*Mtb*) is the human pathogen causing tuberculosis (TB). *Mtb* encodes one RNase HI (RnhC/Rv2228c) and two RNase HII (RnhB/Rv2902c and RnhD/Rv0776c). The non-pathogenic model organism, *Mycobacterium smegmatis*, encodes RnhB, RnhC, and RnhD homologs (MSMEG 2442, MSMEG 4305, and MSMEG 5849) and an additional RNase HI enzyme named RnhA (MSMEG 5562). The activities of the two mycobacterial RNase HI enzymes were characterized in vitro, revealing that they incise within tracts of four or more ribonucleotides in duplex DNA but not an embedded mono-ribonucleotide (20–23). Interestingly, *rnhA* and *rnhC* deletions are synthetically lethal in *M. smegmatis*, showing that RNase HI activity is essential in mycobacteria (21, 24). RnhB and RnhD are presumed to have RNase HII activity based on sequence similarity to the *E. coli* protein. RnhD is capable of hydrolyzing R-loops in vitro and in vivo and its expression complements the temperature-dependent growth defect of the *E. coli* RNase H mutant (25, 26). Deletion of *M. smegmatis rnhB* does not cause a growth defect, impact the level of genomic RNase HII substrates, or increase genome instability (27). Finally, our previous study revealed a division of labor among all *M. smegmatis* RNase H enzymes in tolerance to oxidative stress (21).

Two Mycobacterial RNase Hs, RnhC and RnhD, are bifunctional proteins. RnhD encodes, in addition to the RNase HII domain, a (p)ppGpp Synthetase domain, and both modules are inactive in isolation (25, 26). RnhC is composed of an N terminal RNase HI domain and an autonomous C-terminal acid phosphatase domain, homologous to CobC, a α-ribazole phosphatase involved in vitamin B12 biosynthesis (22, 23). Deletion of *rnhC* in *M. smegmatis* results in diminished vitamin B12 levels in cells cultured in carbon limited acidic medium (28). The acid phosphatase activity of RnhC does not play a role in R-loop degradation or the synthetic lethality between *rnhA* and *rnhC* (21, 28).

Here, we investigate the relative contributions of *M. smegmatis* RNase H enzymes on growth, genome stability and antibiotic sensitivity. We find that RNase HII activity is not essential for optimal growth or to maintain genome stability, but overexpression of either *rnhB* or *rnhD* is highly toxic for the bacterium. We show that the deletion of *rnhC*, but not other *rnh* genes, increases the sensitivity of *M. smegmatis* to rifampicin. However, we demonstrate that this higher rifampicin sensitivity is due to a defect in the acid phosphatase activity of RnhC, rather than its RNase H activity. Finally, we show that the inactivation of the acid phosphatase activity of RnhC renders *M. smegmatis* more sensitive to oxidative stress and impairs light dependent carotenoid pigmentation.

## RESULTS

### Depletion of RNase H2 does not sensitize *M. smegmatis* to hydroxyurea

Our previous data revealed that RNase HI activity (supplied by the RNase HI enzymes encoded by *rnhA* and *rnhC*) is essential for mycobacterial viability (21) but RNase HII activity (supplied by the proteins encoded by *rnhB* and *rnhD*) is not. However, the broader nonessential physiologic roles of RNase HII enzymes in mycobacteria are not understood. Although the Δ*rnhD* strain had a similar growth rate to WT, in-frame deletion of *rnhB* slightly impaired bacterial growth (Figures 1A and B). *rnhB* is the third gene of a putative operon composed by *rplS, lepB* and *rnhB*. Ectopic expression of *rnhB*, controlled by the *rplS* native promoter, in the Δ*rnhB* strain did not restore WT growth rate (Figure 1C), indicating that the growth defect observed in the Δ*rnhB* mutant is not due to *rnhB* inactivation. In addition, we found that Δ*rnhBD, ΔrnhABD*, and Δ*rnhBCD* mutants had a similar growth rate than the single Δ*rnhB* mutant (Figure 1B). These experiments indicate that RNase HII activity is dispensable for optimal growth in *M. smegmatis* in tested conditions, even in absence of one of the two RNases HI.

**Figure 1.**
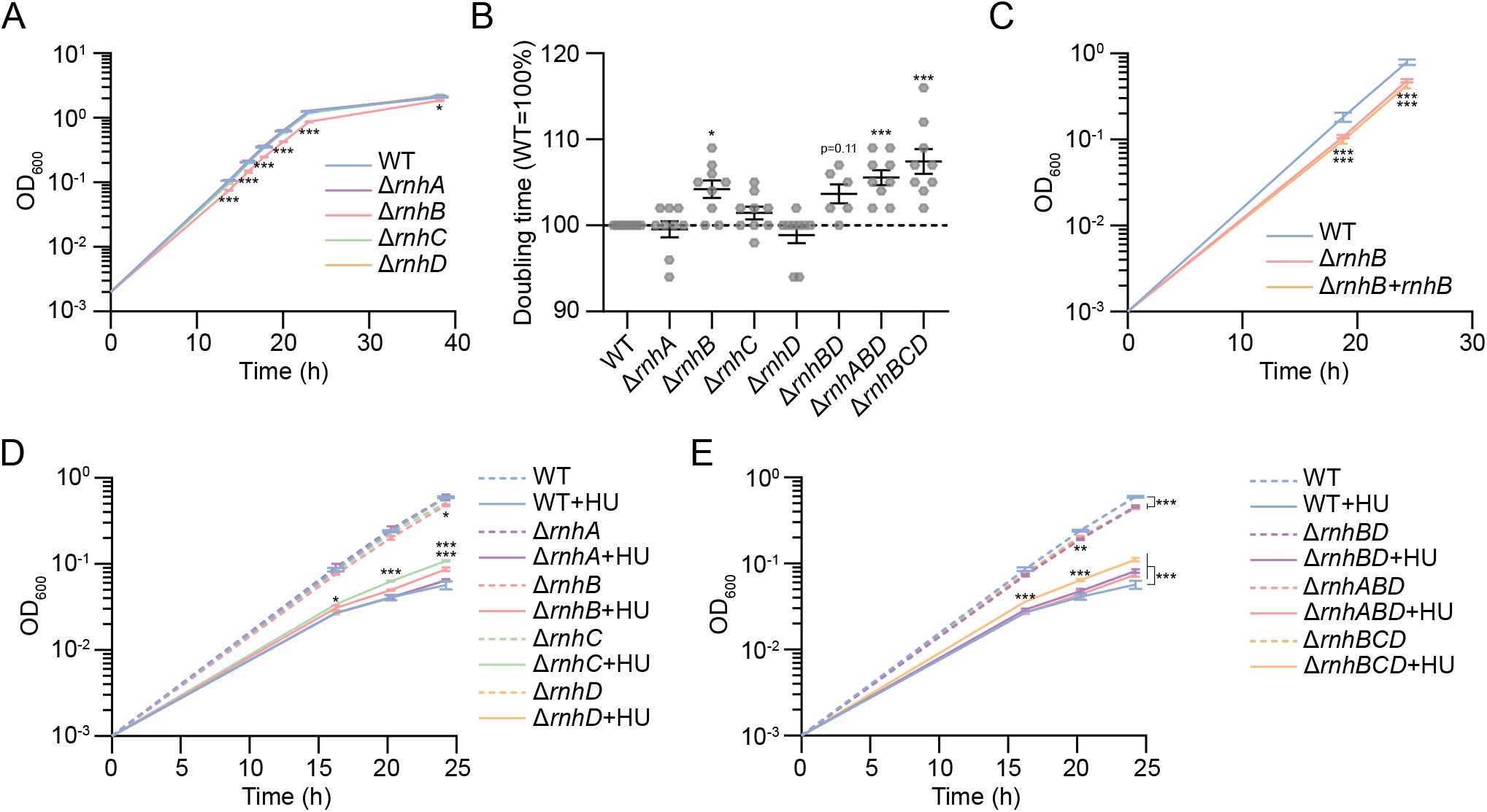
Depletion of RNase HII does not sensitize *M. smegmatis* to hydroxyurea. **(A)**, **(C)**, **(D)**, and **(E)** Bacterial growth curves of indicated strains. In **(C)**, *rnhB* is expressed ectopically under its native promoter (*rpls* upstream sequence). In **(E)**, hydroxyurea (HU) was added to cultures at T0. **(B)** Doubling times of indicated strains cultivated in log phase. Results shown are means (± SEM) of biological triplicates (**A**, **C**, **D**, and **E**) or from biological replicates symbolized by grey dots **(B)**. Stars above or under the means mark a statistical difference with the reference strain (WT) (*, P<0.05; **, P<0.01; ***, P<0.001). p-values were obtained on log-transformed data by two-way (**A**, **C**, **D**, and **E**) or one-way (**B**) ANOVA with a Bonferroni post-test.

Hydroxyurea (HU) is an inhibitor of the ribonucleotide reductase enzyme that increases the rNTP/dNTP ratio and may enhance misincorporation of ribonucleotides into DNA by the replicative polymerase, a genomic lesion that is excised by RNase HII in many systems (29). To determine whether RNase HII activity defends against HU toxicity, we measured the effect of HU treatment on WT and *rnh* mutant growth. We found that HU treatment caused a similar growth defect in WT, Δ*rnhA*, and Δ*rnhD* strains (Figure 1D). Surprisingly, the growth defect caused by HU was slightly reduced in Δ*rnhB* and Δ*rnhC* mutants (Figure 1D and E).

### Overexpression of *rnhB* and *rnhD* is toxic

The slightly better tolerance of the Δ*rnhB* and Δ*rnhC* strain to HU could suggest that RNase HII processing of genomic ribonucleotides could be detrimental in the setting to HU treatment. To test the effect of enhanced RNase HII activity, we measured the impact of RNase H overexpression (OE) on growth. We constructed multi-copy plasmids with *rnhA, rnhB, rnhC*, or *rnhD* under the control of the strong *groEL* promoter. Whereas we obtained many hyg^R^ colonies after transformation with empty, *rnhA* OE, *rnhC* OE, or *rnhD* OE plasmids, no colonies were obtained from the *rnhB* OE plasmid transformation (Figure S1), suggesting that high level of *rnhB* expression is lethal for *M. smegmatis*.

To quantify the lethality of RNase H OE, we expressed *rnhA, rnhB, rnhC* or *rnhD* from an Anhydrotetracycline (ATc) inducible promoter (tet promoter). In absence of inducer, the growth rate of all strains was similar (Figure 2A). Addition of ATc did not impact the growth of the control strain (empty vector) and OE of *rnhA* or *rnhC* caused a weak growth defect. However, OE of RNase HII genes strongly impacted bacterial growth, with the strongest effect observed with *rnhB* overexpression (Figure 2A). Despite minimal effect in liquid culture, growth inhibition by *rnhA* OE was observed on agar medium supplemented with ATc (Figure 2B). As observed in liquid culture, OE of *rnhB* and *rnhD*, but not *rnhC*, caused a strong growth defect on solid medium. Together, these results show that OE of multiple RNase Hs can inhibit *M. smegmatis* growth and that enhanced RNase HII activity is highly toxic.

**Figure 2.**
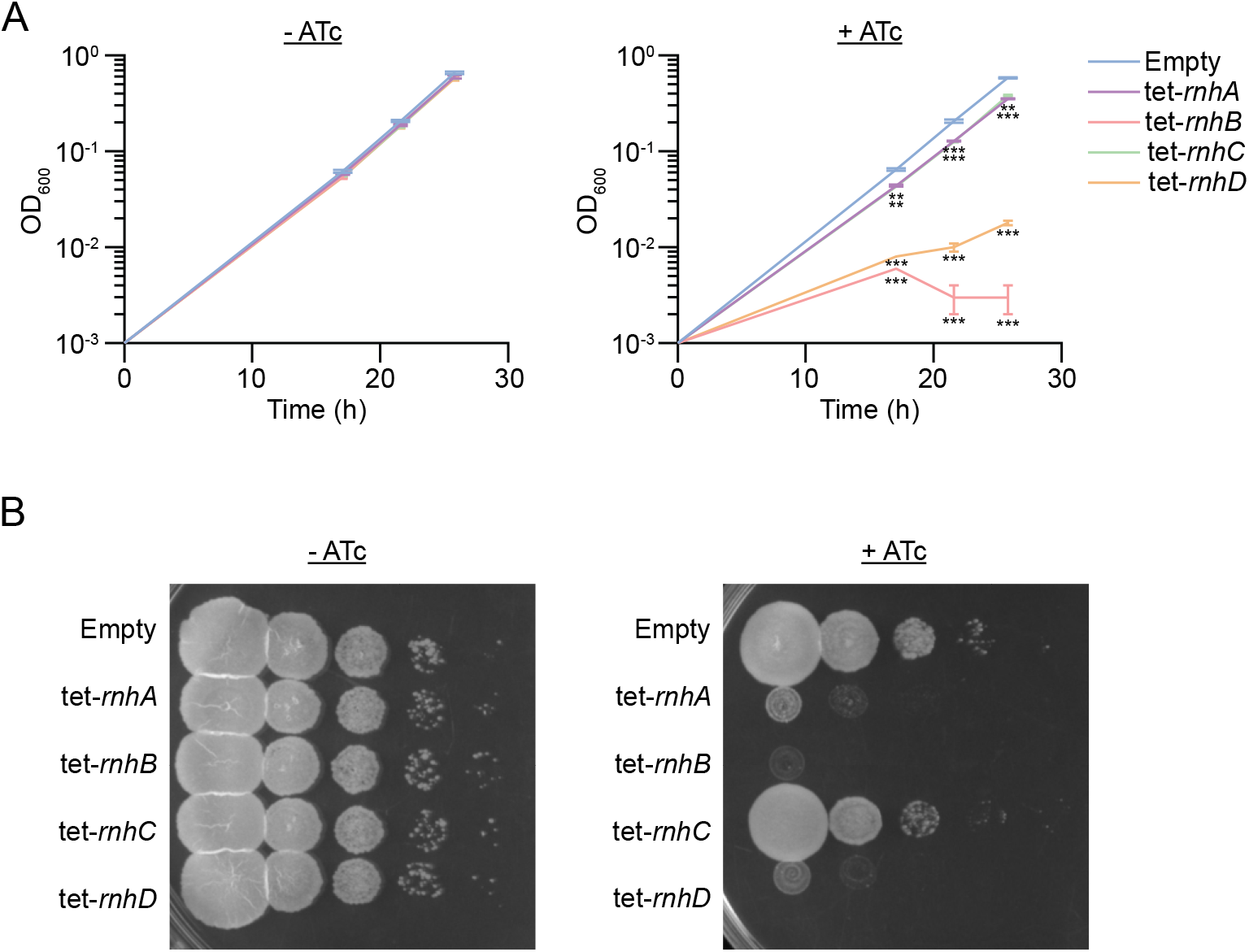
Overexpression of mycobacterial RNase HII inhibits mycobacterial growth. **(A)** Bacterial growth curves of strains carrying an empty vector or an inducible (tet=Anhydrotetracycline (ATc) inducible promoter) *rnhA, rnhB, rnhC*, or *rnhD* in absence or in presence of inducer. Results shown are means (± SEM) of biological triplicates. Stars under the means mark a statistical difference with the reference strain (empty vector) (*, P<0.05; **, P<0.01; ***, P<0.001). p-values were obtained on log-transformed data by two-way ANOVA with a Bonferroni post-test. **(B)** Growth of 10 fold dilutions of the indicated strains on agar medium in absence or presence of inducer. Pictures are representative of experiments performed in triplicate.

### Depletion of Mycobacterial RNase HII does not affect mutation frequency

RNase HII activity is anti-mutagenic in *Bacillus subtilis* but not in *E. coli* (13, 15). To study the impact of RNase H deletion on mutagenesis in mycobacteria, we measured the frequency of rifampicin resistance (rif^R^), conferred by substitution mutations in the rifampin resistance determining region (RRDR) of the *rpoB* gene, in WT and *rnh* mutants. We found similar spontaneous rif^R^ frequencies in WT, Δ*rnhBD, ΔrnhABD*, and Δ*rnhBCD* strains (figure 3A). By sequencing RRDR of rif^R^ colonies, we compared the mutation spectrum of WT and *rnhBD*. In WT, we detected a majority of A>G or T>C and G>A or C>T mutations with a minority of other mutations (Figure 3A), and no discernable difference was observed between strains, indicating no effect of the RNase H system on base substitution mutagenesis.

**Figure 3.**
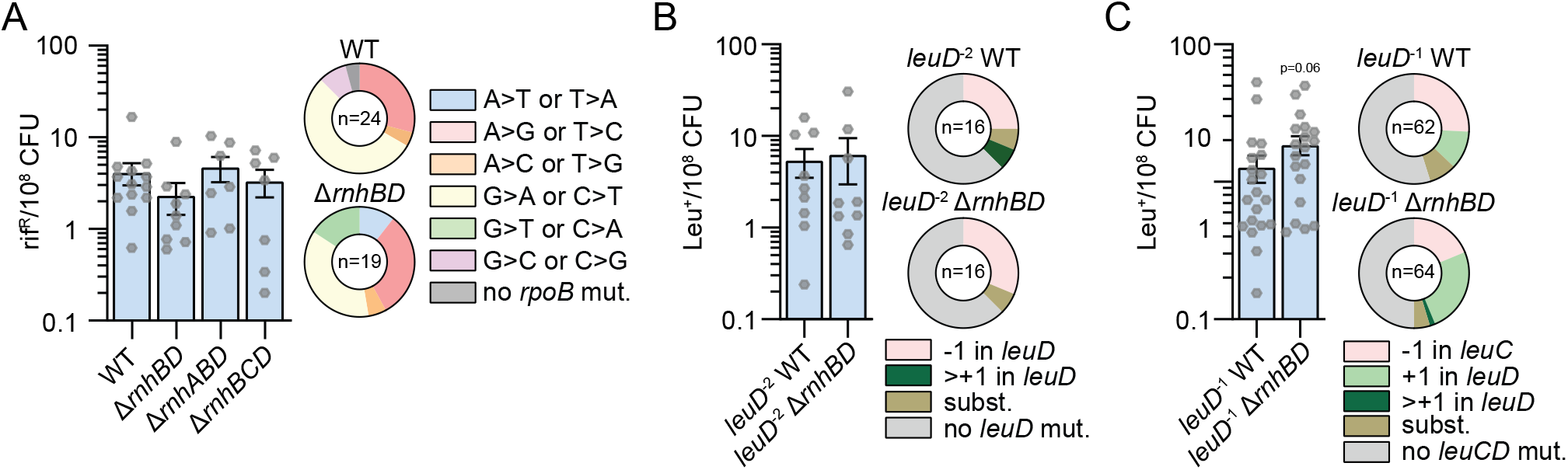
Depletion of Mycobacterial RNase H does not impact substitution or frameshift mutagenesis. **(A)** Rifampicin resistance frequency (rif^R^) or **(B,C)** leucine prototrophy frequency (leu+) in indicated strains. Strains carry a 2-base pair (*leuD*^-2^) **(B)** or a 1-base pair (*leuD*^-1^) **(C)** deletion in the second codon of *leuD* conferring leucine auxotrophy. Results shown are means (± SEM) of data obtained from biological replicates symbolized by grey dots. Stars above bars mark a statistical difference with the reference strain (WT, leuD^-1^ WT, or leuD^-2^) (*, P<0.05). p-values were obtained on log-transformed data by one-way ANOVA with a Bonferroni post-test. Pie charts shows relative frequencies of nucleotide changes (symbolized by color) detected in *rpoB* of rif^r^ in the indicated strains **(A)** or *leuC/leuD* of leu+ in the indicated strains **(B,C)**. The number of sequenced rif^R^ or leu+ is given in the center of each pie chart.

In eukaryotes, deletion of Rnase HII causes frameshift mutations (FS) due to excision of embedded genomic ribonucleotides by topoisomerase (30, 31). FS mutagenesis is not detectable by measuring rif^R^ frequency because RpoB is essential for viability and therefore frameshift mutations would be lethal. To measure the impact of mycobacterial Rnase HII deletion on FS mutagenesis, we used a reporter system, developed in a previous study (32), in which the chromosomal *leuD* gene carries a 1- or 2-base pair deletion in the second codon (*leuD*^-1^ or *leuD*^-2^), conferring leucine auxotrophy. Reversion of either of these mutations by frameshifting confers leucine prototrophy (leu^+^), which is selected on leucine free media. Using the *leuD*^-1^ reporter, similar leu+ frequencies were obtained in WT and Δ*rnhBD* (Figure 3B). In WT, sequencing of *leuD* in leu+ revertants showed that 25% had a −1 FS in *leuD* restoring a functional open reading frame, 6% had a substitution mutation generating a new in-frame start codon, 6% had >+1 bp insertions, and the remainder had no *leuD* mutations (Figure 3B). We observed a weak induction (1.5-fold) of the leu+ frequency in the Δ*rnhBD* mutant using the *leuD*^-1^ reporter (Figure 3C). In WT, 25% of leu+ had a −1 FS localized at the 3’ end of the upstream *leuC* gene, generating a functional in frame LeuC-LeuD fusion, 8% had a substitution mutation generating a new in-frame start codon, 11% had a +1 FS in *leuD*, and 62% had no *leuC* or *leuD* mutation. The Δ*rnhBD* mutant showed 2-fold more +1 FS in *leuD* than WT (Figure 3C). Together, these results show that deletion of mycobacterial RNase HII does not affect the frequencies of spontaneous substitution mutation or −1 FS frequencies and weakly enhances +1 FS events.

### Deletion of *rnhC* confers sensitivity to rifampicin

Our prior data indicated that loss of mycobacterial RNase H enzymes confers sensitivity to oxidative damage (21). Because antibiotics are a source of oxidative stress (33, 34), we investigated the impact of mycobacterial RNase H deletion on antibiotic sensitivity. We tested different classes of antibiotics: rifampicin, streptomycin, ciprofloxacin, and isoniazid which respectively inhibit transcription, translation, DNA replication, and cell wall synthesis. Using a disc diffusion assay, we found that *rnhA* or *rnhD* deletion did not affect the *M. smegmatis* sensitivity to these four antibiotics (Figures 4A, S2B, S2D, and S2E). However, the Δ*rnhB* mutant was more sensitive to rifampicin (Figure 4A), ciprofloxacin (Figure S2B), and streptomycin (Figures S2A and S2D). Additional deletion of other RNase H encoding genes did not exacerbate these phenotypes (Figures 4A, S2B and S2D). The higher sensitivity to streptomycin was detected in two *rnhB* in-frame mutants, constructed independently, showing that the phenotype is due to the *rnhB* deletion and not a random genomic mutation in another gene (Figure S2A). However, ectopic expression of *rnhB* in the Δ*rnhB* strain did not revert the mutant sensitivity to rifampicin and ciprofloxacin (Figures 4B and S2C), indicating that these phenotypes are likely due to a polar effect. Intriguingly, the *rnhB* deletion induced a lower sensitivity to isoniazid (Figures S2E and S2F) which was also not complemented by ectopic expression of *rnhB* (Figure S2G).

**Figure 4.**
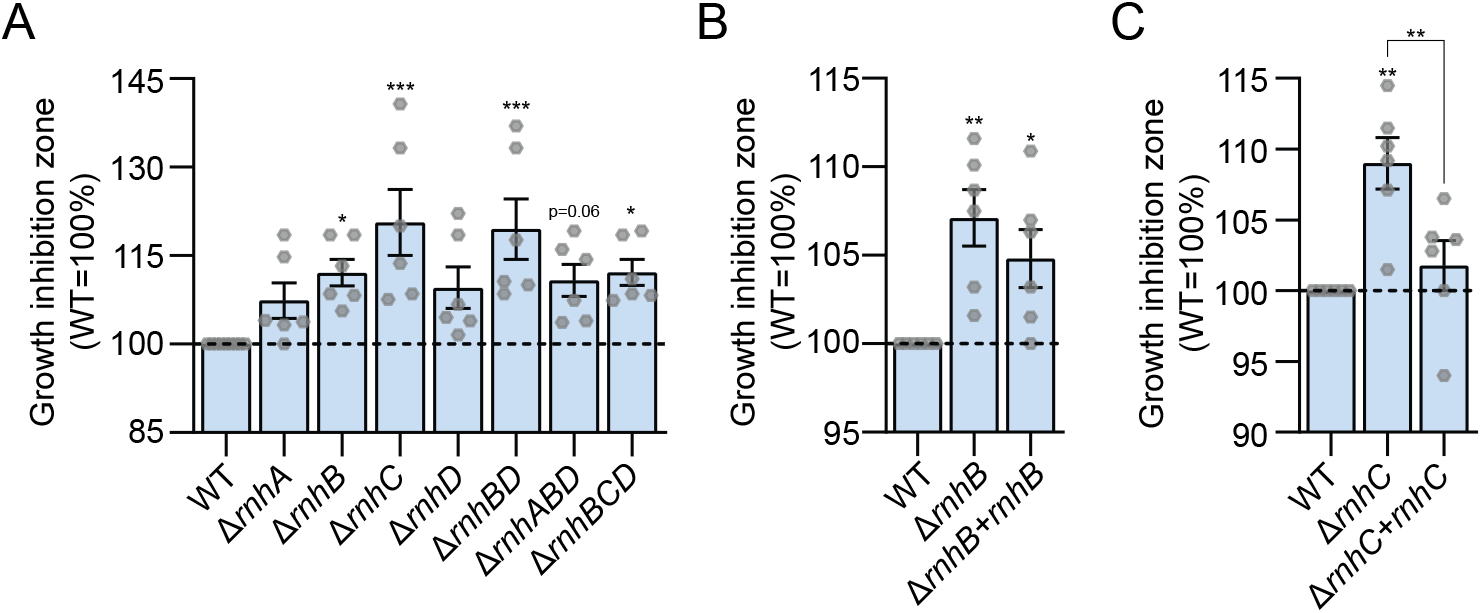
Loss of *rnhC* sensitizes *M. smegmatis* to rifampicin. **(A)**, **(B)**, and **(C)** Sensitivities of indicated strains to rifampicin measured by disc diffusion assay. In **(B)** and **(C)**, *rnhB* and *rnhC* are expressed ectopically under their native promoter (upstream sequences of *rplS* for *rnhB* or of MSMEG_4307 for *rnhC*). Results shown are means (± SEM) of data obtained from biological replicates symbolized by grey dots. Stars above the means mark a statistical difference with the reference strain (WT **(A)**, Δ*rnhB* **(B)**, or Δ*rnhC* **(C)**) (*, P<0.05; **, P<0.01; ***, P<0.001). p-values were obtained on log-transformed data by one-way ANOVA with a Bonferroni post-test.

We detected a higher sensitivity of the Δ*rnhC* mutant to rifampicin by disc diffusion (Figure 4A), but not ciprofloxacin (Figure S2B), streptomycin (Figure S2D), or isoniazid (Figure S2E), which was not exacerbated by the deletion of other RNase H enzymes. A higher sensitivity of the Δ*rnhC* mutant to rifampicin was also detected by microplate resazurin assay (Figure S3A) as well as on agar medium containing rifampicin (Figure S3B). Δ*rnhC* sensitivity was detected in two independent *rnhC* in-frame mutants (Figure S3C) and ectopic expression of *rnhC* in the Δ*rnhC* mutant restored the WT sensitivity (Figures 4C and S3D) indicating that RnhC confers rifampicin tolerance. This finding confirms recently published results using an *M. smegmatis rnhC* mutant that was hypersensitive to Rifampin using an MIC based assay (35).

### Rifampicin sensitivity of the Δ*rnhC* mutant is not due to RNase HI activity

RnhC is a bifunctional protein composed by an N-terminal RNase HI domain and a C-terminal acid phosphatase domain (22). These domains function autonomously in vitro, but the in vivo substrates of the C terminal acid phosphatase domain are not known. To determine which enzymatic activity of the RnhC protein is involved in rifampicin tolerance, we complemented the Δ*rnhC* mutant with *rnhC* alleles encoding catalytic dead mutants of the RNase HI activity (*rnhC*^D73N^) or the acid phosphatase activity (*rnhC*^H173A^) (21, 22). Ectopic expression of *rnhC*^D73N^ in the Δ*rnhC* mutant restored rifampicin sensitivity to the same degree as wild type *rnhC* (Figure 5A), indicating that a defect in its RNase HI activity is not the cause of the higher sensitivity of the Δ*rnhC* mutant. However, ectopic expression of *rnhC*^H173A^, which does not have acid phosphatase activity but retains RNase H activity (22), in the Δ*rnhC* strain did not restore rifampicin tolerance (Figure 5A). We obtained a similar result by plating serial dilutions of these strains on agar supplemented with 5μg/ml of rifampicin (Figure 5B). Importantly, our prior data indicates that ectopic expression of the acid phosphatase defective RnhC (*rnhC*^H173A^) in a Δ*rnhAC* double mutant restores viability, showing that the mutated protein is expressed and that the RNase HI activity is intact (21). These results reveal that RnhC acid phosphatase activity protects *M. smegmatis* against rifampicin.

**Figure 5.**
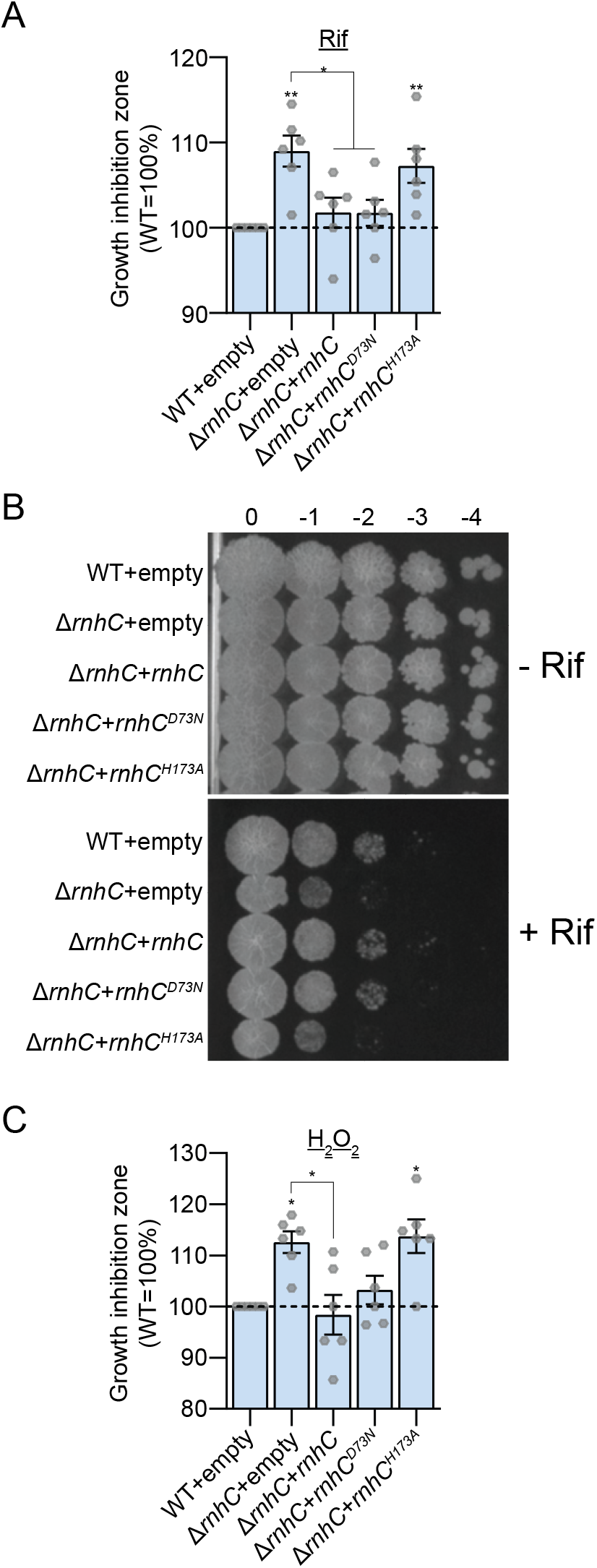
Rifampicin sensitivity of the *rnhC* mutant is due to a loss of the RnhC acid phosphatase activity. Sensitivities of indicated strains (*rnhC*=WT *rnhC, rnhC*^D73N^=RNase HI catalytic mutant of *rnhC, rnhC*^H173A^=acid phosphatase catalytic mutant of *rnhC*) to **(A)** rifampicin and **(C)** H_2_O_2_ measured by disc diffusion assay. WT or mutated versions of *rnhC* are expressed ectopically under their native promoter (upstream sequence of MSMEG_4307). Results shown are means (± SEM) of data obtained from biological replicates symbolized by grey dots. Stars above the means mark a statistical difference with the reference strain (Δ*rnhC*+empty) (*, P<0.05; **, P<0.01; ***, P<0.001). p-values were obtained on log-transformed data by one-way ANOVA with a Bonferroni post-test. **(B)** Growth of indicated strains on agar medium in absence or presence of rifampicin. Pictures are representative of experiments performed in triplicate.

To investigate if the acid phosphatase activity of RnhC could protect bacteria against oxidative stress that accompanies antibiotic treatment, we measured the sensitivity of *rnhC* mutants to H_2_O_2_. We found that the Δ*rnhC* mutant was more sensitive than WT to H_2_O_2_ (Figure 5C). Ectopic expression of *rnhC* or *rnhC*^D73N^ in Δ*rnhC* restored the WT sensitivity to H_2_O_2_, showing that RnhC protects against oxidative stress, but not through its RNase HI activity (Figure 5C). However, complementation of the Δ*rnhC* mutant by the acid phosphatase defective RnhC (*rnhC*^H173A^) did not restore rifampicin tolerance, showing that the acid phosphatase activity of RnhC protects *M. smegmatis* against oxidative stress.

### Phosphatase activity of RnhC is involved in light-dependent pigmentation of *M. smegmatis* colonies

When exposed to light, *M. smegmatis* displays weak photochromogenicity which can be observed as an orange yellow pigment (36, 37). After 3 days of incubation in the dark, colonies of both WT and Δ*rnhC* were white (Figures 6A). WT colonies became yellow when agar plates were kept three days at room temperature in the light (Figures 6A, B, and C). We observed that the Δ*rnhC* mutant had a defect of yellow pigmentation after three days of light exposure (Figures 6A, B and, C), a phenotype which was not visible 11 days later (Figure 6C). However, deletion of *rnhA, rnhB* and, *rnhD*, alone or in combination, did not alter colony pigmentation (Figures 6A and C). The pigmentation defect of the Δ*rnhC* mutant was successfully complemented by expression of either WT *rnhC* or *rnhC*^D73N^, but not by the *rnhC*^H173A^ (Figure 6B), revealing that phosphatase activity, but not RNase HI activity, of RnhC is involved in light-inducible pigmentation of *M. smegmatis*.

**Figure 6.**
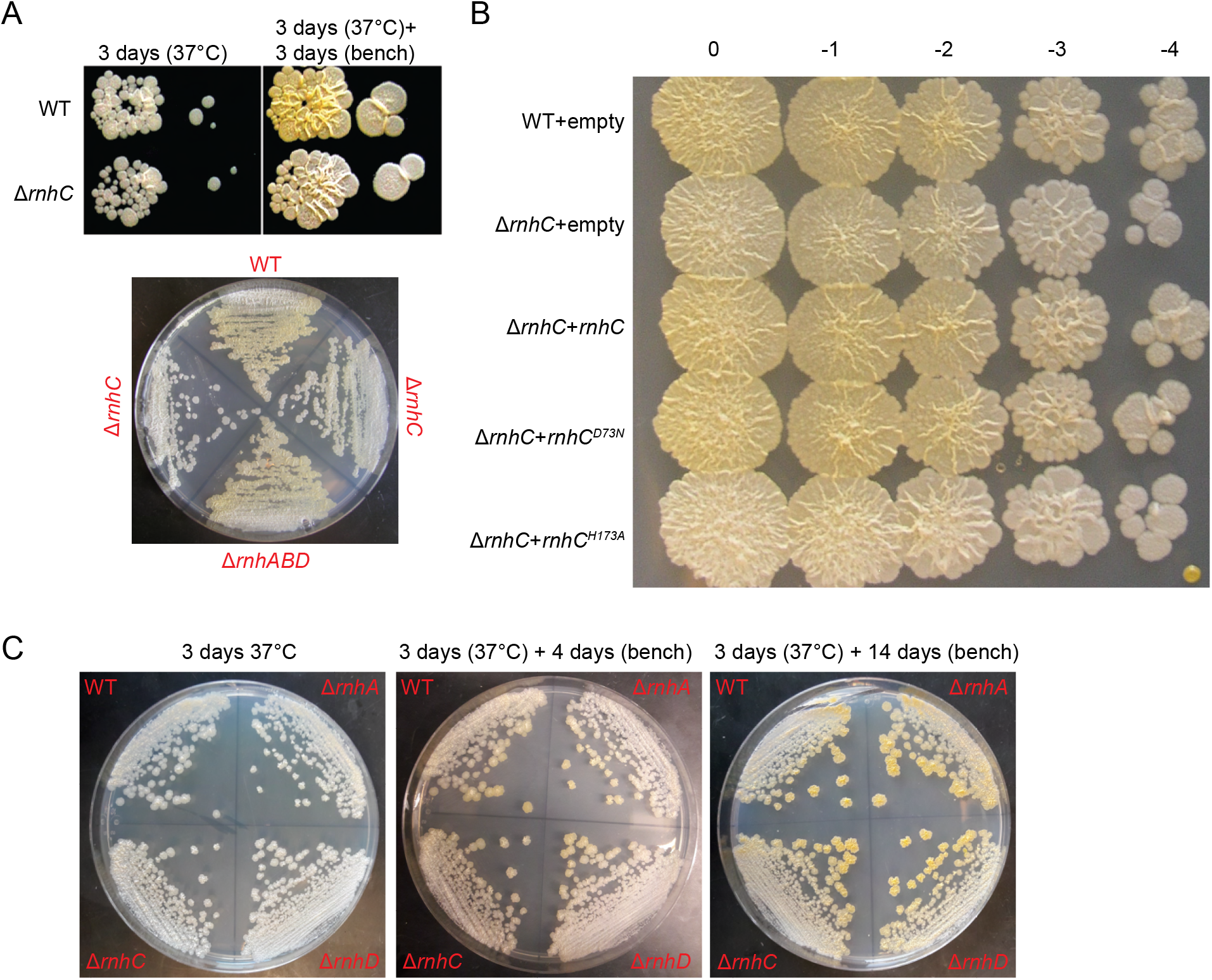
Acid Phosphatase activity of RnhC is involved in the light-dependent pigmentation of *M. smegmatis*. **(A), (B)** and **(C)** Colony pigmentation of indicated strains (*rnhC*=WT *rnhC, rnhC*^D73N^=RNase HI catalytic mutant of *rnhC, rnhC*^H173A^=acid phosphatase catalytic mutant of *rnhC*) cultivated on agar medium in indicated time and conditions of incubation. WT or mutated versions of *rnhC* are expressed ectopically under their native promoter (upstream sequence of MSMEG_4307).

## DISCUSSION

### Enigmatic biological functions of mycobacterial RNase HII

Unlike in *E. coli* and *Bacillus subtilis*, RNase HI activity is essential for the viability of *M. smegmatis* (21, 24, 38, 39). Although biologic basis for this phenotype is still unclear, a defect in processing lagging-strand RNA primers or R-loop degradation is suspected. In this study, we have interrogated the roles of RNase HII activity in mycobacteria. Loss of the two *M. smegmatis* RNase HII enzymes did not affect bacterial viability or mutagenesis in contrast to findings in other bacteria (13, 15). However, we demonstrate that overexpression of RNase HII (either encoded by *rnhB* or *rnhD*) strongly inhibits *M. smegmatis* growth.

The main biological function of RNase HII is excision of embedded genomic ribonucleotides misincoporated by the replicative DNA polymerase (16–18). The absence of phenotype of the *M. smegmatis* RNase HII mutant could be explained by three possibilities: 1) The genomic incorporation of ribonucleotides is rare in mycobacteria or is condition specific, possibly in nonreplicating states during which the rNTP:dNTP may increase; 2) Genomic ribonucleotides are present but are better tolerated by mycobacteria due to more flexible DNA polymerases or more efficient DNA repair pathways; 3) Alternative systems have redundant activities with mycobacterial RNase HII. Minias *et al*. showed that alkaline hydrolysis of DNA is detectable at similar levels in WT and Δ*rnhB*, indicating that ribonucleotides are incorporated in mycobacterial genomes (27). The extreme toxicity of RNase HII overexpression, which we postulate may be due to uncontrolled genomic ribonucleotide excision, may suggest that model 2 is operative and that under physiologic conditions RNase HII activity is restrained to prevent genotoxicity. Further experimentation will be needed to understand the role of RNase HII in mycobacteria.

### RnhC protects against rifampicin through its acid phosphatase activity

The RNase HI enzyme encoded by *rnhC*, while coessential with RnhA, also contains an autonomous C terminal acid phosphatase domain that does not contribute to its essential function (21, 22). Al-Zubaidi *et al*. recently demonstrated that the *rnhC* mutant of *M. smegmatis* led to the accumulation of R-loops and a sensitivity to rifampicin (35), a finding (rif sensitivity) we replicate in this study. However, our data indicates that the RNase HI activity of the RnhC protein is not required for the rifampicin resistance phenotype and that the acid phosphatase activity is the important biochemical activity. These data suggest that the accumulation of R-loops, while clearly shown to be due to loss of *rnhC* (35) and likely due to a defect of the RNase HI activity of RnhC, may not be linked to rifampicin sensitivity.

### RnhC controls colony pigmentation of *M. smegmatis*

In this work, we reveal that the acid phosphatase domain of RnhC is involved in light-dependent yellow pigmentation of colonies in *M. smegmatis*. This pigmentation is known to be due to the synthesis of carotenoids, controlled by the SigF sigma factor (36, 37). In *M. smegmatis*, as well as in other orange-pigmented mycobacteria, the carotenoid molecule involved in colony pigmentation is isorenieratene (36, 40–43). Isorenieratene production is catalyzed by five metabolic steps in *M. smegmatis*, involving CrtE, CrtB, CrtI, CrtY, and CrtU enzymes (36). In the absence of SigF, *crt* operon is not expressed and pigmentation is abolished (36, 37).

The mechanism by which RnhC controls *M. smegmatis* colony pigmentation is unknown. The acid phosphatase domain of RnhC is homologous with the CobC α-ribazole phosphatase, involved in vitamin B12 biosynthesis (22, 23). Vitamin B12 can be synthesized de novo by some prokaryotes, thought complex metabolic routes, involving several dozens of enzymes (44). CobC is involved in the last step of the Vitamin B12 synthesis, catalyzing dephosphorylation of adenosylcobalamin-5’-phosphate (45). One possibility is that RnhC could be involved in the dephosphorylation of some isorenieratene precursors, but the ultimate mechanism will require further investigation.

### Model for the interrelationship of rifampin sensitivity, oxidative stress, and pigment production

We show that C-terminal acid phosphatase domain of RnhC confers tolerance to rifampicin and H_2_O_2_, and is required for carotenoid biosynthesis. Previous studies demonstrate that production of carotenoids protects *M. smegmatis* against oxidative stress (36, 37). The bactericidal effect of some antibiotics, including rifampicin, has also been linked to oxidative stress (33, 34). One model to understand the higher sensitivity of the Δ*rnhC* mutant to rifampicin is that a failure to dampen antibiotic induced oxidative stress due to the absence of carotenoids enhances the potency of the antimicrobial. However, we cannot exclude the possibility that RnhC is involved in the synthesis of other unidentified components protecting *M. smegmatis* from rifampicin.

The bifunctional RnhC protein is conserved in several mycobacteria, including pathogenic species *Mtb* and *Mycobacterium leprae* (22, 23, 28). Rifampicin is a cornerstone antibiotic used to treat *Mtb*. As proposed by Al-Zubaidi *et al*. (35), The higher sensitivity of the *M. smegmatis* Δ*rnhC* mutant to rifampicin opens the possibility of targeting RnhC to potentiate rifampicin and reduce the effective dose or duration of rifampicin treatment as well as attenuate the frequency of resistance acquisition. However, based on our data, compounds that sensitize to rifampicin should not be optimized based on the RNase HI activity but instead the acid phosphatase activity of RnhC. Unlike *M. smegmatis* or *M. marinum, Mtb* is assumed to not synthesize carotenoids even though it encodes carotenoid cleavage oxygenases (46). Therefore, the mechanisms proposed here may differ in *M. tuberculosis* and future studies will be needed to assess the link between RnhC, carotenoid production, and rifampicin sensitivity in *Mtb*.

## Methods

### Bacterial strains

Strains of this work are listed in Supplementary Table 1. *Escherichia coli* and *M. smegmatis* strains were cultivated at 37°C in, respectively, Luria-Bertani (LB) medium and Middlebrook 7H9 medium (Difco) supplemented with 0.5% glycerol, 0.5% dextrose, 0.1% Tween 80. Streptomycin and hygromycin were respectively used at 5 μg/ml and 50 μg/ml.

### Plasmids and deletion mutants construct

Plasmids of this study were constructed in *E. coli* DH5α and are listed in Supplementary Table 2. Complementation plasmids were constructing by cloning ORFs together with their 5’ flanking regions (~500bp), amplified by PCR using *M. smegmatis* mc^2^155 genomic DNA as template and primers listed in Supplementary Table 3, into integrative pDP60 vector digested with EcoR1. *rnh* inducible and *rnh* overexpression plasmids were constructed by cloning *rnhA, rnhB, rnhC*, or *rnhD* ORFs, amplified by PCR using *M. smegmatis* mc^2^155 genomic DNA as template and primers listed in Supplementary Table 3, into pmsg419 digested with *ClaI* and pmv261 digested with BamHI, respectively. Markerless and in-frame gene deletions were performed as described in Barkan et al., 2011 using pAJF067 derivatives containing ~500-bp regions flanking the gene to be deleted. ORF-flanking fragments were amplified by PCR using *M. smegmatis* mc^2^ 155 genomic DNA as template and primers listed in Supplementary Table 3 and were cloned into pAJF067 digested with NdeI. All cloning were performed using In-Fusion recombination-based cloning method (Takara) and plasmids were introduced into *M. smegmatis* strains by electroporation.

### Growth

7H9 medium was inoculated with log growth phase bacteria to OD_600_=0.001. Growth was measured by monitoring OD_600_ for two days. For *rnh*-inducible experiments, log growth phase bacteria cultured without inducer (Anhydrotetracycline: ATc) were back diluted in fresh 7H9 medium supplemented with 50 nM of inducer to OD_600_=0.001. For growth on agar medium, serial dilutions of log growth phase bacteria cultivated without inducer were spotter (5 μl) on 7H10 medium supplemented with 50 nM of ATc and incubated at 37°C for 72 h.

### Disc diffusion assay

Log growth phase bacteria were diluted in 3 ml of pre-warmed top agar (7H9, 6 mg/ml agar) to an OD_600_ of 0.01 and plated on 7H10. A filter disc was put on the dried top agar and was spotted with 2.5μl of 100 mg/ml rifampicin (rif), 100 mg/ml streptomycin (sm), 10 mg/ml ciprofloxacin (cip), 10 mg/ml isoniazid (INH), or 10M H_2_O_2_. The diameter of the growth inhibition zone was measured after incubation for 48h at 37°C.

### Agar-based assay

Log growth phase bacterial cultures were diluted to an OD_600_ of 0.1. 5 μl of serial dilutions (10^0^ to 10^-5^) were spotted on 7H10 or 7H10 supplemented 5-10 μg/ml rifampicin or 100-200 μg/ml streptomycin. Pictures were taken after 3 days incubation at 37°C.

### Resazurin assay

Log growth phase bacterial cultures were diluted to an OD_600_ of 0.0005 in 7H9 supplemented with various concentrations of antibiotics. Cultures were incubated 2 days at 37°C into 96 wells plates (100 μl of culture per plate), sealed with parafilm. 30 μl of 0.2 mg/ml resazurin was added to cultures and incubated 24h at 37°C.

### Mutation frequency

For measurement of substitution mutation frequency, bacteria were grown to log phase in 7H9 medium from a single colony, back-diluted at an OD_600_ of 0.001 in fresh medium and cultured for 24 h. Cells (OD_600_ ~0.5) were concentrated 40-fold by centrifugation/pellet resuspension and 100 μl of a 10^-6^ dilution was plated on 7H10 agar whereas 200 μl was plated on 7H10 supplemented with 100 μg/ml rifampicin. For the measurement of FS mutation frequency, cells were cultivated in 7H9 medium supplemented with 50 μg/ml leucine. 40-fold concentrated cultures were plated on 7H10 supplemented with 50 μg/ml leucine (100 μl of a 10^-6^ dilution) or 7H10 (200 μl of a 10^0^ dilution). The number of independent cultures used to measure the mutation frequency is indicated by the number of grey dots in each bar of graphs. Mutation spectrum was established by sequencing, using primers listed in Supplementary Table 3, the rifampicin resistance determining region (RRDR) of the *rpoB* gene of isolated rif^R^ colonies or *leuCD* genes of isolated leu^+^ colonies, amplified by PCR. For each strain, sequenced colonies were picked among at least six biological replicates.

### Colony pigmentation assay

Log phase bacteria cultivated in 7H9 were plated (serial dilution spotting or streaking of an undiluted culture) on 7H10 and incubated in dark at 37°C for 3 days. Plates were kept at room temperature for 3 days or more under natural light.

## Supporting information

Merged SI file

## Data availability

Further information and requests for resources and reagents should be directed to and will be fulfilled by Dr. Michael Glickman. Plasmids and strains generated in this study will be made available on request. All data generated in this study are presented in the Figures and Tables. Any additional information required to reanalyze the data reported in this paper is available from the lead contact upon request.

## Acknowledgments

This work is supported by NIH (NIH grant #AI064693) and this research was funded in part through the NIH/NCI Cancer Center Support Grant P30CA008748. P. Dupuy was supported in part by a « Jeune Scientifique » salary award from the French National Institute of Agronomic Science (INRA). We thank all Glickman lab members for helpful discussions.

## Author contributions

P.D. and M.G. designed research; P.D. performed research; P.D. and M.G. analyzed data; P.D. and M.G. wrote the paper.

## Supplementary data

Supplementary Figures S1-S3 and Supplementary Tables S1-S3.

