## Supplementary material for "The C-terminal acid phosphatase module of the RNase HI enzyme RnhC controls rifampicin sensitivity and light-dependent colony pigmentation of *Mycobacterium smegmatis*": Merged SI file

### Supplementary information

#### Supplementary figure legends

**Figure S1. Transformation efficiency of plasmids carrying *rnh* genes under constitutive strong promoter.** Transformation of 50 ng of an empty vector conferring the resistance to rifampicin or plasmids carrying *M. smegmatis rnh* genes under *groEL* promoter selected on agar medium supplemented with hygromycin. Upper pictures: 1/10 of transformation mixtures incubated 6 days at 37 °C on 7H10 medium supplemented with hygromycin. Lower pictures: 10 µl spots of serial dilutions of transformation mixtures after 3 days incubation at 37 °C on 7H10.

**Figure S2. *rnh* mutants sensitivities to antibiotics.** . **(A)** Growth of indicated strains on agar medium in presence of indicated concentrations of streptomycin. **(B)**, **(C)**, **(D)**, **(E)**, and **(G)** Sensitivities of indicated strains to indicated antibiotics (Cipro=ciprofloxacin, Sm=streptomycin, INH=isoniazid) measured by disc diffusion assay. Results shown are means (± SEM) of data obtained from biological replicates symbolized by grey dots. Stars above the means mark a statistical difference with the reference strain (WT) (*, P<0.05; **, P<0.01; ***, P<0.001). p-values were obtained on log-transformed data by one-way ANOVA with a Bonferroni post-test. **(F)** Picture of a representative biological replicate shown in **(E)** graph.

**Figure S3. Δ*rnhC* sensitivity to rifampicin.** **(A)** and **(C)** Growth of indicated strains in 7H9 supplemented with indicated concentrations of rifampicin and resazurin blue dye (redox indicator). Resazurin turns pink by aerobic respiration of metabolically active cells. **(B)** and **(D)** Growth of indicated strains on agar medium in absence or presence of rifampicin (**(B)** 10 µg/ml, **(D)** 5 µg/ml). Pictures are representative of experiments performed in triplicate.

### Supplementary tables

| Strains | Genetic background | reference or source |
|---|---|---|
| *Mycobacterium smegmatis* | | |
| *M. smeg* MC$^2$155 | WT | (Snapper et al., 1990) |
| Mgm4087 | *rnhA*::hyg | (Gupta et al., 2017) |
| Mgm4085 | Δ*rnhB* | (Gupta et al., 2017) |
| PDS121 | Δ*rnhB* | This work |
| Mgm4084 | Δ*rnhC* | (Gupta et al., 2017) |
| PDS123 | Δ*rnhC* | This work |
| Mgm4089 | *rnhD*::zeo | (Gupta et al., 2017) |
| Mgm4090 | Δ*rnhB*, *rnhD*::*zeo* | (Gupta et al., 2017) |
| PDS143 | Δ*rnhA*, Δ*rnhB, rnhD::zeo* | This work |
| PDS145 | Δ*rnhB*, Δ*rnhC, rnhD::zeo* | This work |
| PDS622 | *leuD*$^{-1}$ | (Dupuy et al., 2022) |
| PDS626 | Δ*rnhB*, *rnhD*::*zeo*, *leuD*$^{-1}$ | This work |
| PDS630 | *leuD*$^{-2}$ | (Dupuy et al., 2022) |
| PDS634 | Δ*rnhB*, *rnhD*::*zeo*, *leuD*$^{-2}$ | This work |
| *Escherichia coli* | | |
| DH5α | F$^-$ Φ80*lac*ZΔM15 Δ(*lac*ZYA-*arg*F) U169 *rec*A1 *end*A1 *hsd*R17($r_k^-$, $m_k^+$) *pho*A *sup*E44 *thi*-1 *gyr*A96 *rel*A1 λ$^-$ | Lab collection |

Table S1: strains used in this study

| Plasmids | description | Cloning enzyme sites | Cloning primers (infusion reaction) | References or sources |
|---|---|---|---|---|
| pAJF067 | Gene replacement vector ($Hyg^R$, *galK*, *sacB*) | | | (Fay and Glickman, 2014) |
| pDB60 | Mycob. integr. vector ($Strep^R$, attP(L5)) | | | Lab Stock |
| pmsg419 | ATc-on system vector ($hyg^R$, OriMyc) | | | Lab Stock |
| pmV261 | expression vector GroEL strong promoter ($hyg^R$, OriMyc) | | | (Stover et al., 1991) |
| pDP3 | pAJF067 derivative-rnhA deletion | *NdeI* | ODP79-ODP80 +ODP81-ODP82 | This work |
| pDP4 | pAJF067 derivative-rnhB deletion | *NdeI* | ODP13-ODP14 +ODP15-ODP16 | This work |
| pDP5 | pAJF067 derivative-rnhC deletion | *NdeI* | ODP17-ODP18 +ODP19-ODP20 | This work |
| pDP6 | pAJF067 derivative-rnhD deletion | *NdeI* | ODP21-ODP22 +ODP23-ODP24 | This work |
| pDP104 | pAJF067 $leuD^{-1}$ frameshift | | | (Dupuy et al., 2022) |
| pDP105 | pAJF067 $leuD^{-2}$ frameshift | | | (Dupuy et al., 2022) |
| pDP18 | pMV261-hyg derivative-*rnhA* overexpression | *BamHI* | ODP123-ODP124 | This work |
| pDP19 | pMV261-hyg derivative-*rnhB* overexpression | *BamHI* | ODP125-ODP126 | This work |
| pDP20 | pMV261-hyg derivative-*rnhC* overexpression | *BamHI* | ODP127-ODP128 | This work |
| pDP21 | pMV261-hyg derivative-*rnhD* overexpression | *BamHI* | ODP129-ODP130 | This work |
| pDP41 | pmsg419 derivative-$rnhA^{streptag}$ overexpression | *ClaI* | ODP224-ODP226 | |
| pDP43 | pmsg419 derivative-$rnhB^{streptag}$ overexpression | *ClaI* | ODP227-ODP229 | |
| pDP45 | pmsg419 derivative-$rnhC^{streptag}$ overexpression | *ClaI* | ODP230-ODP232 | |
| pDP47 | pmsg419 derivative-$rnhD^{streptag}$ overexpression | *ClaI* | ODP233-ODP235 | |
| pDP60 | pDB60 derivative-*rnhB* expression under *rpls* promoter | *EcoRI* | ODP163-ODP164 +ODP165-ODP166 | |
| pRGM54 | pDB60 derivative: *rnhC* expression under MSMEG_4307 promoter | | | (Gupta et al., 2017) |
| pRGM56 | pDB60 derivative: $rnhC^{D73D}$ expression under MSMEG_4307 promoter | | | (Gupta et al., 2017) |
| pRGM55 | pDB60 derivative: $rnhC^{H173A}$ expression under MSMEG_4307 promoter | | | (Gupta et al., 2017) |

Table S2. Plasmids used in this study

| primers | Sequences (5'→3')$^*$ | Targets | vector and cloning sites |
|---|---|---|---|
| Deletion mutants constructs | | | |
| ODP13 | **CTAGTATGCATCATA**ATCCGCTCGGACAACC | fw upstream r*nhB* | pAJF067 (*NdeI*) |
| ODP14 | AATCACGGTTCGAGGC | rev upstream r*nhB* | pAJF067 (*NdeI*) |
| ODP15 | CCTCGAACCGTGATTTGCGCGAAGATGGGC | fw downstream r*nhB* | pAJF067 (*NdeI*) |
| ODP16 | **CTAGGCAATTGCATA**ACGAGCATCTCAAGCCG | rev downstream r*nhB* | pAJF067 (*NdeI*) |
| ODP17 | **CTAGTATGCATCATA**AGATCGACTCGGTGCG | fw upstream r*nhC* | pAJF067 (*NdeI*) |
| ODP18 | CTCGACGAGAACCTTCAC | rev upstream r*nhC* | pAJF067 (*NdeI*) |
| ODP19 | AAGGTTCTCGTCGAGAACCAGACCGCGTACC | fw downstream r*nhC* | pAJF067 (*NdeI*) |
| ODP20 | **CTAGGCAATTGCATA**ATCTGCCGCGGTTGG | rev downstream r*nhC* | pAJF067 (*NdeI*) |
| ODP21 | **CTAGTATGCATCATA**AATCGCTTGCCCAGC | fw upstream r*nhD* | pAJF067 (*NdeI*) |
| ODP22 | GAGGTCGATGCCCACG | rev upstream r*nhD* | pAJF067 (*NdeI*) |
| ODP23 | GTGGGCATCGACCTCAACCGGTTCGAGAAGC | fw downstream r*nhD* | pAJF067 (*NdeI*) |
| ODP24 | **CTAGGCAATTGCATA**ACGCTGATCGCGTTCG | rev downstream r*nhD* | pAJF067 (*NdeI*) |
| ODP79 | **CTAGTATGCATCATA**ACCACCGTCGCAGCC | fw upstream r*nhA* | pAJF067 (*NdeI*) |
| ODP80 | GTGGATGATGACGGGATCC | rev upstream r*nhA* | pAJF067 (*NdeI*) |
| ODP81 | CCCGTCATCATCCACGACGAACTCGCGACCC | fw downstream r*nhA* | pAJF067 (*NdeI*) |
| ODP82 | **CTAGGCAATTGCATA**AGCCGTCGGTGACTTCG | rev downstream r*nhA* | pAJF067 (*NdeI*) |

$^*$15 bp homology with linearized vectors in bold letters.
15 bp homology between two PCR fragments underlined

| | | | |
|---|---|---|---|
| *rnh* constitutive overexpression constructs | | | |
| ODP123 | **AGACAATTGCGGATC**TGGTGATCTCGACGCC | fw *rnhA* | pmv261-hyg (*BamHI*) |
| ODP124 | **TTCTGCAGCTGGATC**TCAACGGAGCGAAGTACCC | rev *rnhA* | pmv261-hyg (*BamHI*) |
| ODP125 | **AGACAATTGCGGATC**AAGGCGCGATTCATCG | fw *rnhB* | pmv261-hyg (*BamHI*) |
| ODP126 | **TTCTGCAGCTGGATC**TCATCCACTGCCCATCTTC | rev *rnhB* | pmv261-hyg (*BamHI*) |
| ODP127 | **AGACAATTGCGGATC**ACGACGACGTGGTGCG | fw *rnhC* | pmv261-hyg (*BamHI*) |
| ODP128 | **TTCTGCAGCTGGATC**CTACAGGTACGCGGTCTGG | rev *rnhC* | pmv261-hyg (*BamHI*) |
| ODP129 | **AGACAATTGCGGATC**AGAACATTCGGAATCGTCACG | fw *rnhD* | pmv261-hyg (*BamHI*) |
| ODP130 | **TTCTGCAGCTGGATC**ATCTGACCGACGCACCC | rev *rnhD* | pmv261-hyg (*BamHI*) |

$^*$15 bp homology with linearized vectors in bold letters.

| | | | |
|---|---|---|---|
| *rnh* inducible overexpression constructs | | | |
| ODP224 | **CAGAAAGGAGGCCAT**GTGAGCCAGGATCCCG | fw *rnhA* | pmsg419 (*ClaI*) |
| ODP226 | **AGGTCGACGGTATCG**CTACTTTTCGAACTGCGGGTGGCTCCAACGGAGCGAAGTACCCG | rev *rnhA* | pmsg419 (*ClaI*) |
| ODP227 | **CAGAAAGGAGGCCAT**GTGCCCGGCTCGTG | fw *rnhB* | pmsg419 (*ClaI*) |
| ODP229 | **AGGTCGACGGTATCG**CTACTTTTCGAACTGCGGGTGGCTCCATCCACTGCCCATCTTCG | rev *rnhB* | pmsg419 (*ClaI*) |
| ODP230 | **CAGAAAGGAGGCCAT**GTGAAGGTTCTCGTCGAGG | fw *rnhC* | pmsg419 (*ClaI*) |
| ODP232 | **AGGTCGACGGTATCG**CTACTTTTCGAACTGCGGGTGGCTCCACAGGTACGCGGTCTGGTT | rev *rnhC* | pmsg419 (*ClaI*) |
| ODP233 | **CAGAAAGGAGGCCAT**ATGCACCACCCCCCGTAC | fw *rnhD* | pmsg419 (*ClaI*) |

| | | | |
|---|---|---|---|
| ODP235 | **AGGTCGACGGTATCG**CTACTTTTCGAACTGCGGGTGGCTCCAGGCCTGCAGCTTCTCGAA | rev *rnhD* | pmsg419 (*ClaI*) |

*15 bp homology with linearized vectors in bold letters.
Streptavidin tag underlined.

*rnhB* complementation construct

| | | | |
|---|---|---|---|
| ODP163 | **TCCAGCTGCAGAATT**AACATCGCGATCGCACTG | fw rplS promoter | pDB60 (*EcoRI*) |
| ODP164 | CGGTGACACTTCCTTG | rev rplS promoter | pDB60 (*EcoRI*) |
| ODP165 | AAGGAAGTGTCACCGGTGCCCGGCTCGTG | fw *rnhB* | pDB60 (*EcoRI*) |
| ODP166 | **GATAAGCTTCGAATT**TCATCCACTGCCCATCTTC | rev *rnhB* | pDB60 (*EcoRI*) |

*15 bp homology with linearized vectors in bold letters.
15 bp homology between two PCR fragments underlined

Table S3. Primers used in this study

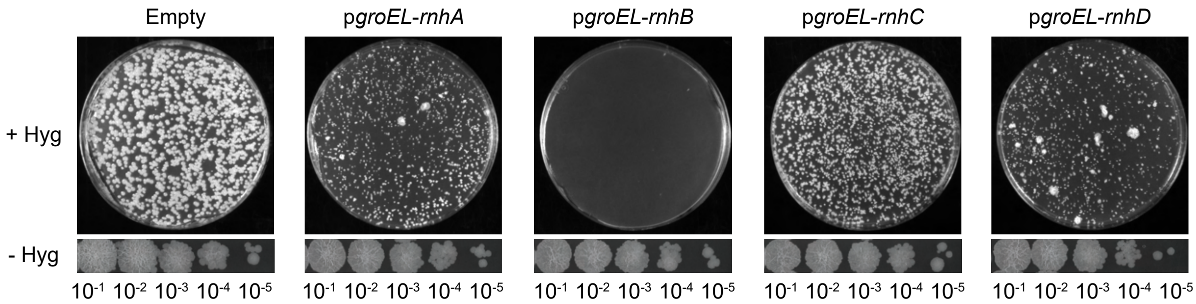


**Figure S1. Transformation efficiency of plasmids carrying *rnh* genes under constitutive strong promoter.** Transformation of an empty vector conferring the resistance to rifampicin or plasmids carrying *M. smegmatis* genes *rnh* genes under *groEL* promoter selected on agar medium supplemented with hygromycin. Upper pictures: 1/10 of transformation mixtures incubated 6 days at 37 °C on 7H10 medium supplemented with hygromycin. Lower pictures: 10 µl spots of serial dilutions of transformation mixtures after 3 days incubation at 37 °C on 7H10.

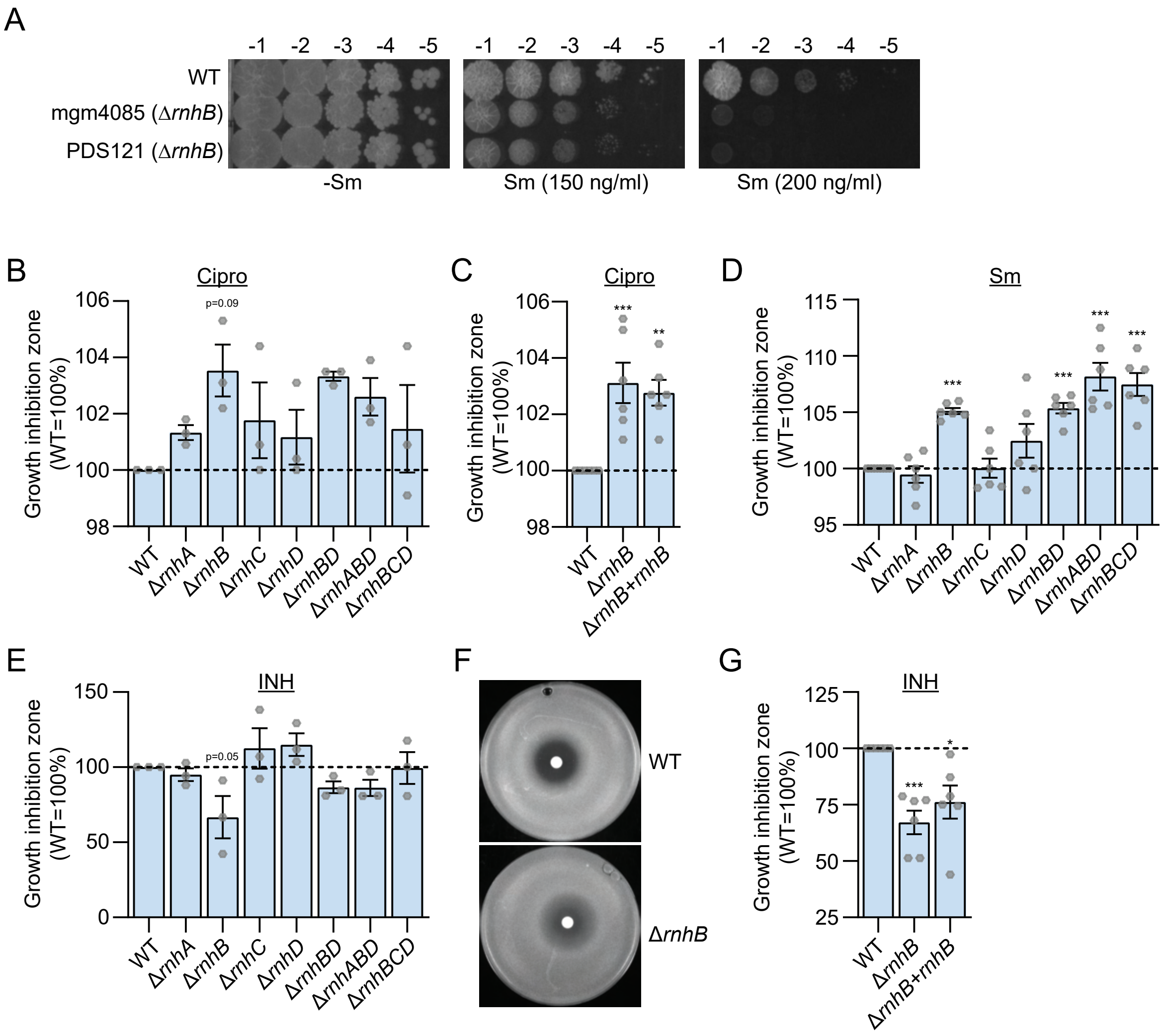


**Figure S2. Antibiotic sensitivity of *rnh* mutant strains.** . **(A)** Growth of indicated strains on agar medium in the presence of indicated concentrations of streptomycin. **(B)**, **(C)**, **(D)**, **(E)**, and **(G)** Sensitivities of indicated strains to indicated antibiotics (Cipro=ciprofloxacin, Sm=streptomycin, INH=isoniazid) measured by disc diffusion assay. Results shown are means (± SEM) of data obtained from biological replicates symbolized by grey dots. Stars above the means mark a statistical difference with the reference strain (WT) (*, P<0.05; **, P<0.01; ***, P<0.001). p-values were obtained on log-transformed data by one-way ANOVA with a Bonferroni post-test. **(F)** Picture of a representative biological replicate from **(E)**.

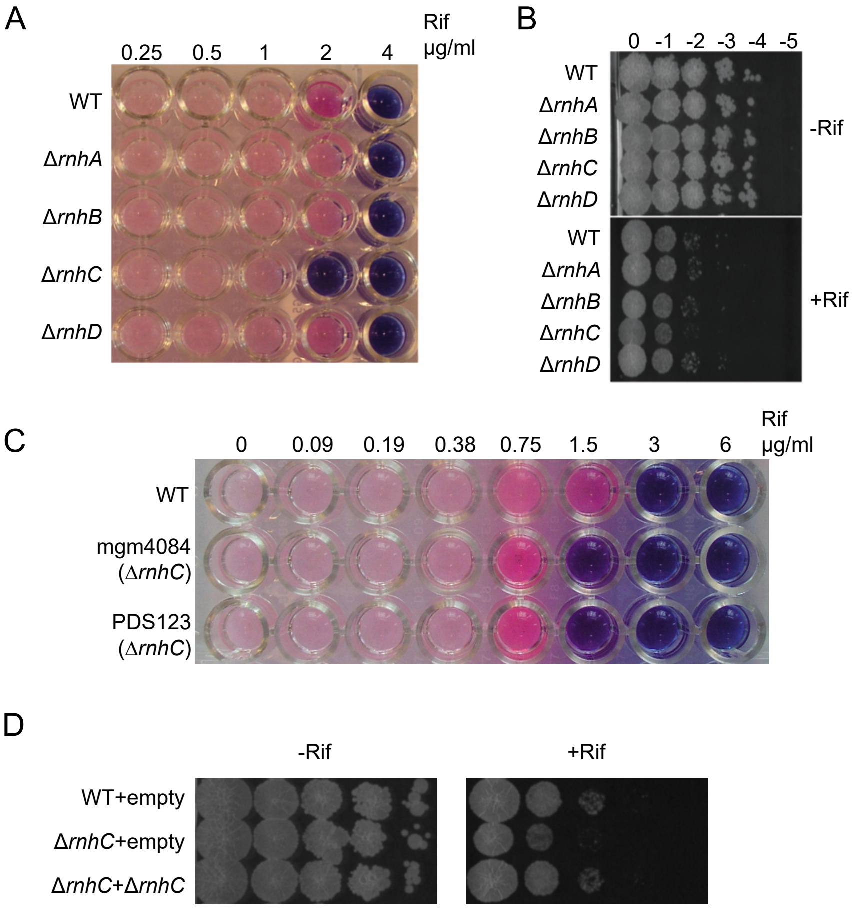


**Figure S3. Δ*rnhC* sensitivity to rifampicin. (A)** and **(C)** Growth of indicated strains in 7H9 supplemented with indicated concentrations of rifampicin and resazurin dye (redox indicator). Resazurin turns pink from aerobic respiration of metabolically active cells. **(B)** and **(D)** Growth of indicated strains on agar medium in absence or presence of rifampicin **((B)** 10 µg/ml, **(D)** 5 µg/ml). Pictures are representative of experiments performed in triplicate.
